# Crowding within synaptic junctions influence the degradation of adenoside nucleotides by CD39 and CD73 ectonucleotidases

**DOI:** 10.1101/2021.09.21.461163

**Authors:** Hadi Rahmaninejad, Tom Pace, Peter Kekenes-Huskey

## Abstract

Synapsed cells can communicate using exocytosed nucleotides like adenosine triphosphate (ATP). Ectonucleotidases localized to a synaptic junction degrade such nucleotides into metabolites like adenosine monophosphate (AMP) or adenosine, oftentimes in a sequential manner. CD39 and CD73 are a representative set of coupled ectonucleotidases, where CD39 first converts ATP and adenosine diphosphate (ADP) into AMP, after which the AMP product is dephosphorylated into adenosine by CD73. Hence, CD39/CD73 help shape cellular responses to extracellular ATP. In a previous study [1] we demonstrated that the rates of coupled CD39/CD73 activity within synapse-like junctions are strongly controlled by the enzymes’ co-localization, their surface charge densities, and the electrostatic potential of the surrounding cell membranes. In this study, we demonstrate that crowders within a synaptic junction, which can include globular proteins like cytokines and membrane-bound proteins, impact coupled CD39/CD73 electronucleotidase activity and in turn, the availability of intrasynapse ATP. Specifically, we simulated a spatially-explicit, reaction-diffusion model for the coupled conversion of ATP→AMP and AMP→adenosine in a model synaptic junction with crowders via the finite element method. Our modeling results suggest that the association rate for ATP to CD39 is strongly influenced by the density of intrasynaptic protein crowders, as increasing crowder density suppressed ATP association kinetics. Much of this suppression can be rationalized based on a loss of configurational entropy. The surface charges of crowders can further influence the association rate, with the surprising result that favorable crowder/nucleotide electrostatic interactions can yield CD39 association rates that are faster than crowder-free configurations. However, attractive crowder/nucleotide interactions decrease the rate and efficiency of adenosine production, which in turn increases the availability of ATP and AMP within the synapse relative to crowder-free configurations. These findings highlight how CD39/CD73 ectonucleotidase activity, electrostatics and crowding within synapses influence the availability of nucleotides for intercellular communication.

## 2 Introduction

Intercellular junctions facilitate cell-to-cell communication[2–4]. Synapsed neurons are a prominent example of cells that form such junctions. Within these junctions, nucleotides such as ATP are commonly used to relay messages between neurons [5]. Ectonucleotidases like CD39 and CD73 rapidly hydrolyze ATP and its metabolites. [6] For this reason, the activity of CD39 and CD73 can strongly influence nucleotide-based communication between synapsed or densely-packed cells, by helping to determine the availability of ATP and AMP to plasma membrane-bound receptors [7].

Recent structural studies suggest that synaptic spaces are typically crowded with biomolecules and scaffolding proteins [8–11]. Examples can include cytokines and other exocytosed cargo, ectodomains of membrane-bound receptors, channels, and pumps, as well as columnar scaffolds tethering juxtaposed synapsed membranes [8, 12–17]. Biochemical processes occurring in crowded environments such as the cell cytosol [18–21], are known to exhibit altered enzyme kinetics, stability and solute transport [22–29] relative to bulk conditions, but have not been robustly examined in synapses. For the cell cytosol, the occupied volume fraction of crowders ranges from 10-50% [30]. Although quantitative estimates for crowding volume fractions in synapses are not readily available, it is reasonable to anticipate that synapses are far from a crowder-free regime. This raises the question, does intrasynapse crowding influence CD39/CD73 activity and if so, how do crowder properties influence enzyme activity?

Many studies have probed enzyme kinetics in crowded media (reviewed in [22]), These studies demonstrate that enzyme kinetics in crowded solutions frequently exhibit attenuated reaction rates given reduced substrate diffusion and enzyme accessibility [28, 29, 31–35]. Further, crowders that have weak interactions with charged substrates can accelerate or hinder diffusion under certain regimes [32, 36]. There are however fewer studies of these processes when they are confined to cellular and organelle membrane surfaces [25, 37]. A recent study evaluated CD39/CD73 activity when the enzymes were confined to narrow and charged synaptic junctions [1] relative to bulk conditions and found that while confinement reduced the ATP association rate to CD39, confinement of the AMP product often increased the production of adenosine. While the study quantified how nucleotidase confinement and electrostatic interactions help shape nucleotide pools within the synapse, the influence of synaptic crowders such as receptors, soluble proteins and scaffolding proteins [12–16], was not examined.

We therefore investigated how crowders modulate the composition of the ATP, AMP, and adenosine nucleotide pool, by controlling CD39/CD73 reaction kinetics (ATP → AMP by CD39, AMP → adenosine by CD73). This was done using a coupled reaction-diffusion model developed in [1] that was adapted to reflect a variety of crowding conditions. Using this model, we quantified how intrasynapse crowders and scaffolding proteins affect 1) CD39/ATP association rates 2) the concerted production of adenosine by CD73, and 3) the availability of ATP and its nucleotides. Our predictions indicate the presence and properties of intrasynaptic crowders can strongly influence CD39 and CD73 kinetics relative to bulk and crowder-free environments, which in turn determine the composition nucleotide pools in the synapse.

## 3 Result and discussion

### 3.1 Steric effects of synaptic crowding on ATP/E1 association kinetics

We first establish that our approach for predicting the ATP/CD39 diffusion-limited association rate recapitulates analytic estimates. The analytic estimate for the association rate, *k*_*on*_, of a substrate to a spherical enzyme in an open (bulk) domain is given by the Kimball-Collins limit [38]

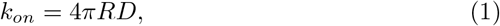

where R is the radius of the CD39 enzyme and D is the diffusion coefficient for the ATP substrate in an uncrowded environment. In Fig. 2 we compare this analytical result to numerical simulations in a large junction approximating bulk conditions. The predicted k_*on,ATP*_ (open square) is within 5% of the analytic estimate (open circle), which validates our modeling approach. In [1] we rationalize that the minor discrepancy can be attributed to our use of a finite-sized simulation domain as opposed to the infinite open domain assumed for the analytic estimate. Within the 8 nm radius synaptic space, the CD39 k_*on,ATP*_ is reduced approximately ten-fold relative to the open domain.

**Figure 1:**
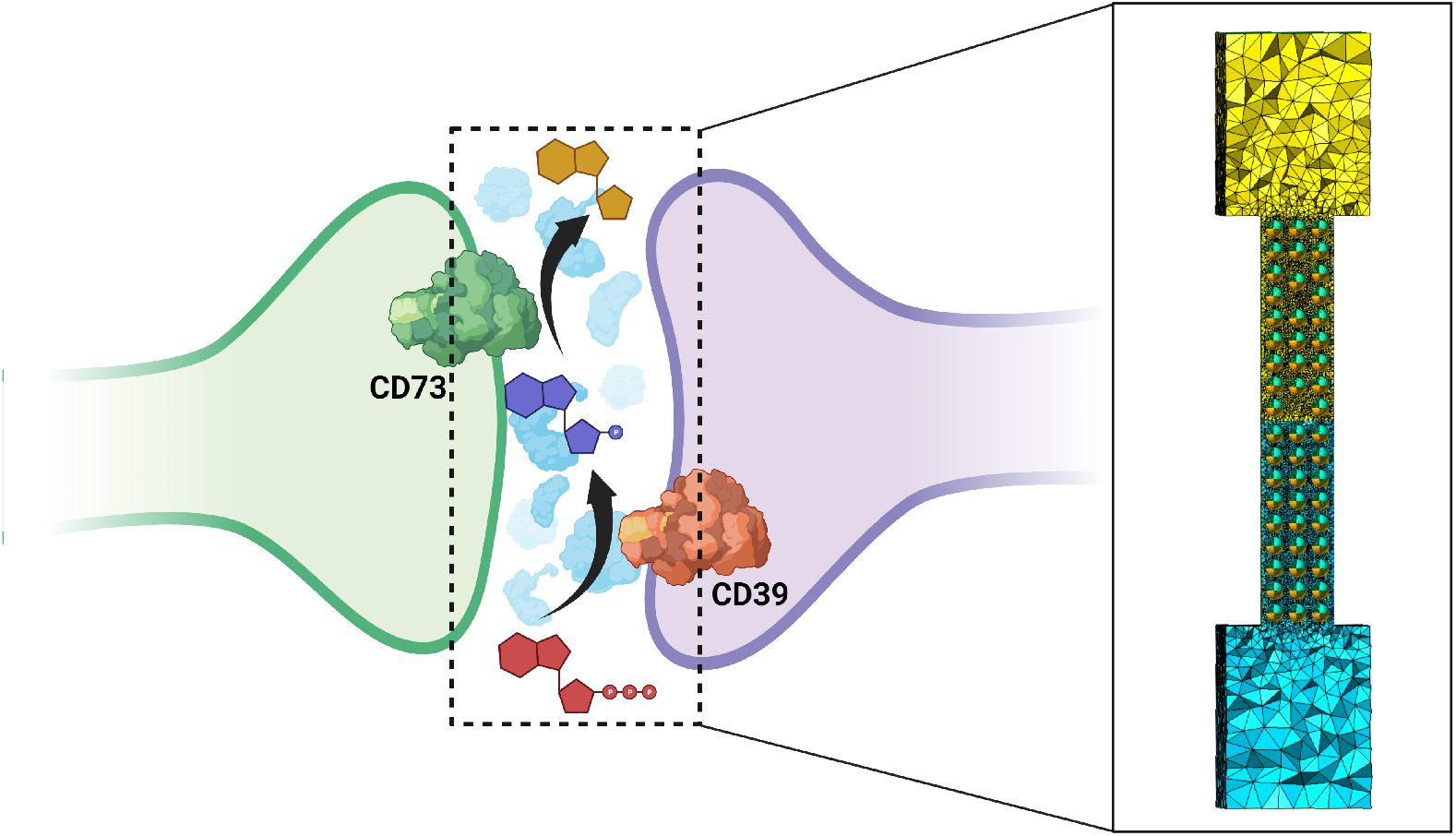
Systems studied in present work. Left) Schematic of a synaptic junction formed between adjacent cells. Coupled ectonucleotidases CD39 and CD73 confined within this space hydrolyze adenosine triphosphate (ATP) into adenosine monophosphate (AMP) and adenosine (Ado) in the presence of other non-reactive crowders. Right) Schematic of the meshed geometry representing the model junction for the Finite Element Analysis. This includes the reservoirs corresponding to the extracellular environment connected to the junction where the spherical crowders are located.

**Figure 2:**
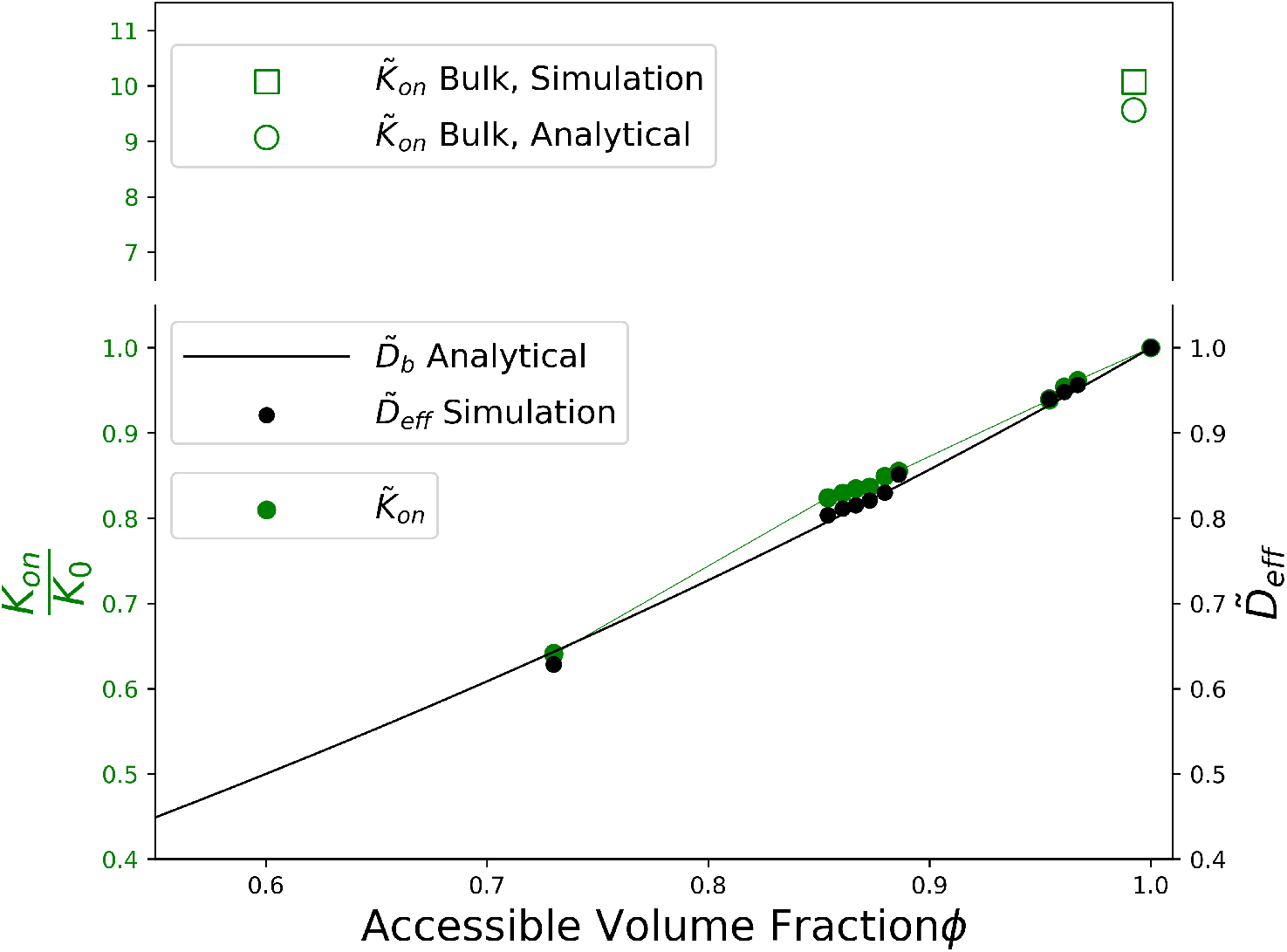
Simulation results under bulk conditions and within a crowded junction. Values for the bulk (uncrowded) condition are shown with hollow symbols, where the analytic result (open circle) is from Eq. 1 for an enzyme radius of 2 nm and the simulation (open square) was conducted with a junction radius of 18.25 nm to approximate bulk conditions. The solid dots and line represent the behavior in a junction of 8 nm radius under varying amounts of crowding, with *ϕ* = 1 representing the uncrowded condition. Black dots are the simulation results for the effective diffusion coefficient normalized to its bulk value. The solid line is the analytical result for the normalized effective diffusion coefficient based on the Hashin-Shtrikman expression in Eq. 2. Green dots are the simulation results for the association rate coefficient normalized the uncrowded condition.

We next introduced uncharged and unreactive (inert) crowders into the synapse to reduce the volume into which ATP can diffuse (*ϕ <* 1). Our predictions for k_*on,ATP*_ are summarized in Fig. 2, which generally show that the association rate decreases as the density of crowders with diameters of 25Å increases. We interpret this trend by noting the dependence of *k*_*on*_ on the effective diffusion coefficient (*D*_eff_) in Eq. 1, where *D* is the diffusion coefficient for ATP in the absence of crowders. Crowders are well-known to reduce ATP diffusion in a manner dependent on the crowder volume fraction. This dependence of a material transport parameter on the packing of spherical inclusions was characterized using an upper bound introduced by Hashin and Shtrikman [39, 40], with the crowders assigned a diffusion coefficient of zero. While Hashin and Shtrikman originally characterized this expression as an upper bound, in practice it is also often a reasonable estimate of the result.

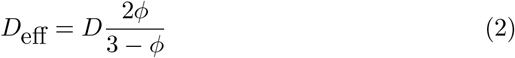

Here to show the effect of accessible volume *ϕ*, we normalized the total flux passing from the pore by its value in the absence of any inclusion (*ϕ* = 0). *D*_eff_ estimates are plotted against the right-hand y-axis of Fig. 2. Here we found that the effective diffusivity is in close agreement with the estimate for *D*_eff_ from Eq. 2, similar to what was reported in [41]. Hence, the primary effect of crowding on the ATP/CD39 association rate can be attributed to hindered diffusion. We report similar findings when the synapse comprises column-like scaffolding proteins, such as the protein leucine-rich glioma inactivated 1 (LGI1) [42] that we represented as cylinders (Fig. S2).

### 3.2 Electrostatic effects of synaptic crowding on ATP/E1 association kinetics

We next evaluated how electrostatic interactions between ATP and charged crowders modulate the CD39/ATP association rate. This was done through solving the Smoluchowski equation (Eq. 9 in Sect. 5.2), using an electrostatic potential estimated via Eq. 10. We have previously validated this approach for predicting the association rate of a substrate to a uniformly charged enzyme in an open domain [43, 44]. Here we extend these results to a charged enzyme embedded in a crowder-free synaptic junction in Fig. 4, where k_*on,ATP*_ is reported for an attractive (V=25mV), neutral, and repulsive potential(V=-25mV), respectively and as a function of normalized reactive surface area, *γ*. Our results for a uniformly reactive sphere are signified by *γ* = 1. The reported values are normalized to the k_*on,ATP*_ obtained for the neutral enzyme and confirm the intuitive result that attractive ATP/CD39 interactions yield association rates faster than the neutral case, and conversely, reduced rates for repulsive interactions. CD39 has a positively-charged active site [45], therefore these intuitive results suggest that its charge would promote association of the ATP anion.

Although neutral crowders intuitively reduced the predicted association rates to CD39, our previous study [32] indicated that crowders with charges complementary to diffusing ATP can enhance diffusion relative to uncharged crowders. We therefore investigated if attractive ATP/crowder interactions would similarly enhance k_*on,ATP*_ İn Fig. 3, we predict k_*on,ATP*_ for positive (attractive) and negative (repulsive) crowders. Our data demonstrate that repulsive interactions reduce k_*on,ATP*_ values relative to neutral crowders. Conversely, association rates in the presence of attractive crowders are faster than both neutral and repulsive crowders. Although the k_*on,ATP*_ values are smaller than the rates predicted for crowder-free conditions under most free volume fractions, a narrow range of volume fractions yield k_*on,ATP*_ values that exceed the crowder-free configuration (see Fig. 3 for the three-dimensional case, and Fig. S2 for the two-dimensional case). Hence, although confinement dramatically reduces association rates to CD39 relative to bulk, this effect can be offset to some degree by attractive crowder/ATP interactions.

**Figure 3:**
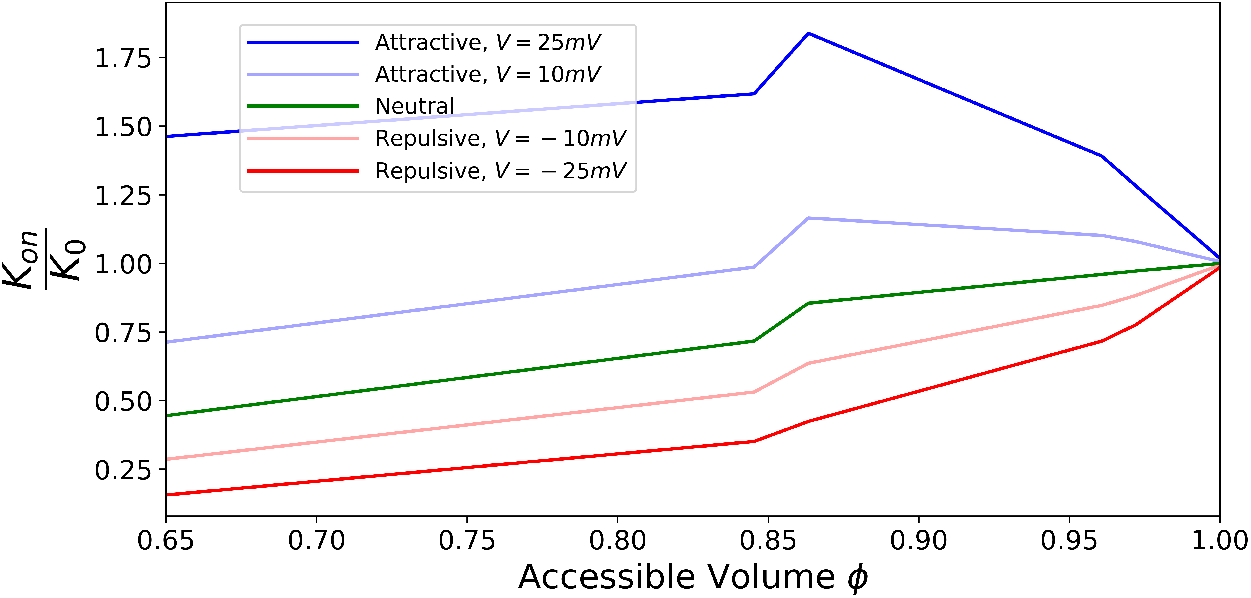
Effect of charged non-reactive crowders on the CD39/ATP reaction rate. As the volume occupied by crowders increases, the free volume fraction (*ϕ*) decreases. Five different electric potential boundary conditions were assigned to the crowders: one neutral condition (zero charge), two potentials that are attractive to ATP, and two potentials that are repulsive to ATP. ATP association rate coefficients are normalized to the crowder-free case (*ϕ* = 1). Normalized association rates above 1 therefore represent higher association rates than the crowder-free condition. This is the case for a select range of free volume fractions, which is wider for the higher attractive potential.

To ease interpretation of how interactions between the substrate and crowders modulate reaction kinetics for the enzyme, we next utilized an approximation from Alsallaq *et al* [46, 47] to estimate a mean-field interaction potential of ATP with CD39. This will be used later to quantify the thermodynamic impact crowders impose on the CD39 association rate. The Alsallaq approximation relates an observed association, *k*_*on*_ rate as a function of its intrinsic association rate, *k*_*on,unch*_ and a centrosymmetric electric potential *U*_*elec*_.

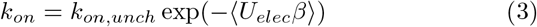

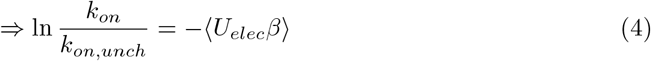

The expression − ⟨*U*_*elec*_*β*⟩ represents the electrostatic energy averaged over a reactive volume, normalized by kT (*β* ≡ *kT* ^*−*1^). This approximation assumes that reacting entities have a narrow reaction region and a centrosymmetric, long-ranged potential [46, 47]. We demonstrate in Fig. 4 that the predicted rates approach estimates based on Eq. 3 as the reactive surface area *γ* → 0.1. Namely, our methodology predicts an association rate within 3% for repulsive interactions versus 14% for attractive interactions. These data establish the reliability of our approach in predicting diffusion-limited association rates and estimating ATP/CD39 interaction potentials governing their association for small reaction patches. Importantly, the reasonable agreement of our modeling results with estimates from Eq. 3 suggests that the equation properly describes the influence of electrostatic interactions on ATP diffusion. Below, we apply this same equation to the influence of other interactions that can be described by a mean force potential.

**Figure 4:**
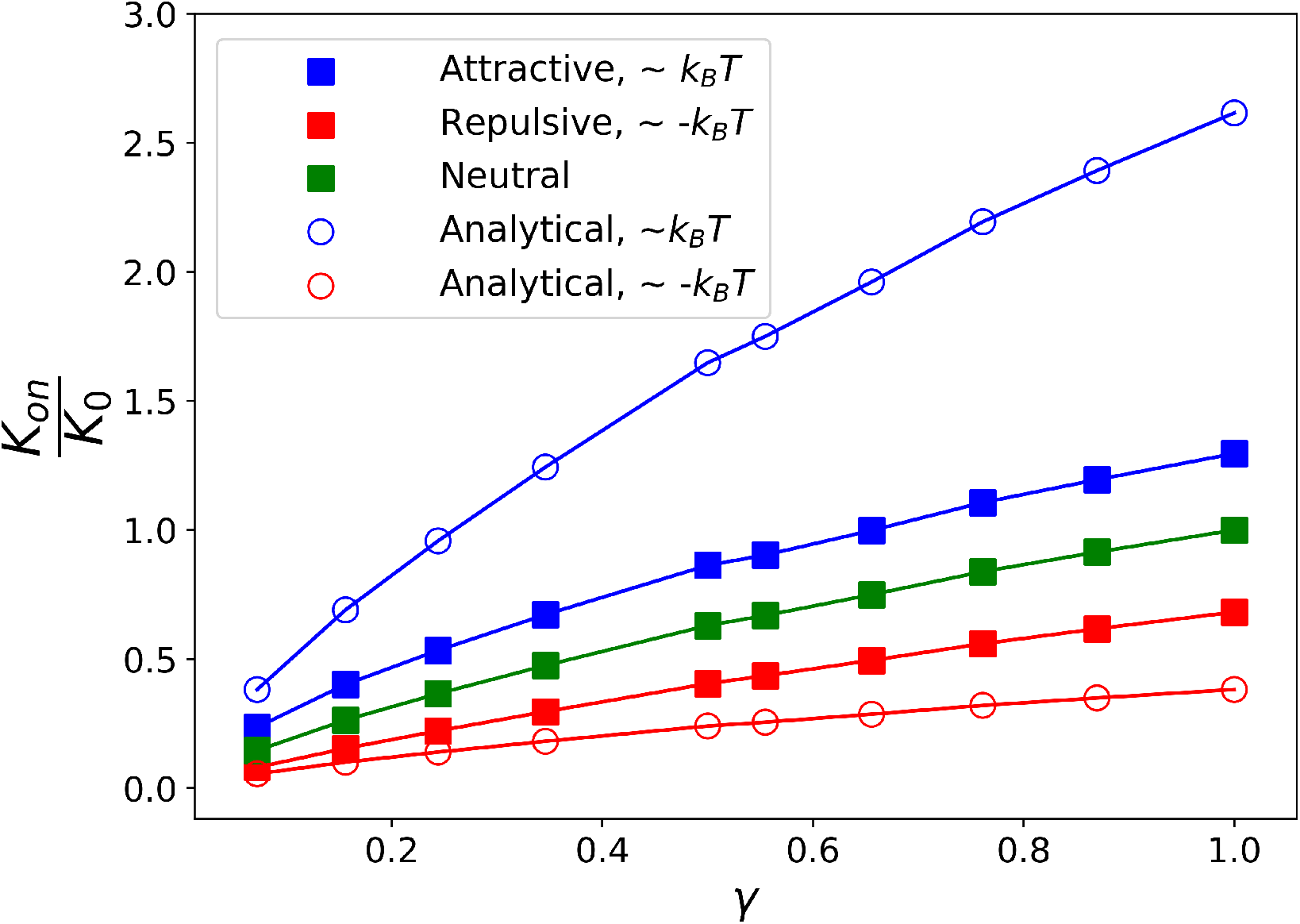
Diffusion-limited ATP/CD39 association rate as a function of the relative reactive patch size *γ. γ* = 1 signifies a uniformly reactive enzyme. The attractive and repulsive refers to the enzyme electrostatic potential with a magnitude of 25*mV*. The analytical results are obtained based on Eq. 3, which is only valid at small reactivity region on the enzyme. All of the data are normalized to the neutral value of an enzyme which is uniformly reactive.

### 3.3 Energetic basis for crowder-mediated CD39/ATP association kinetics

Our results so far indicate that the volume excluded by crowders significantly reduces diffusion and subsequently, substrate binding kinetics. We therefore used Eq. 3 and results from the previous section to express the effect of inert crowders as an effective potential. Our data in Fig. 5 indicate that the exponentiated effective potential, which is obtained by Eq. 4 is a nearly linear function of free volume fraction, *ϕ*. In the absence of crowders, U=0, while U is repulsive (*U* > 0) for increasing crowder densities (See panel b in Fig. 5). These data indicate that the effect of crowders can be expressed as an effective potential, U that modulates the ATP/CD39 association rates.

**Figure 5:**
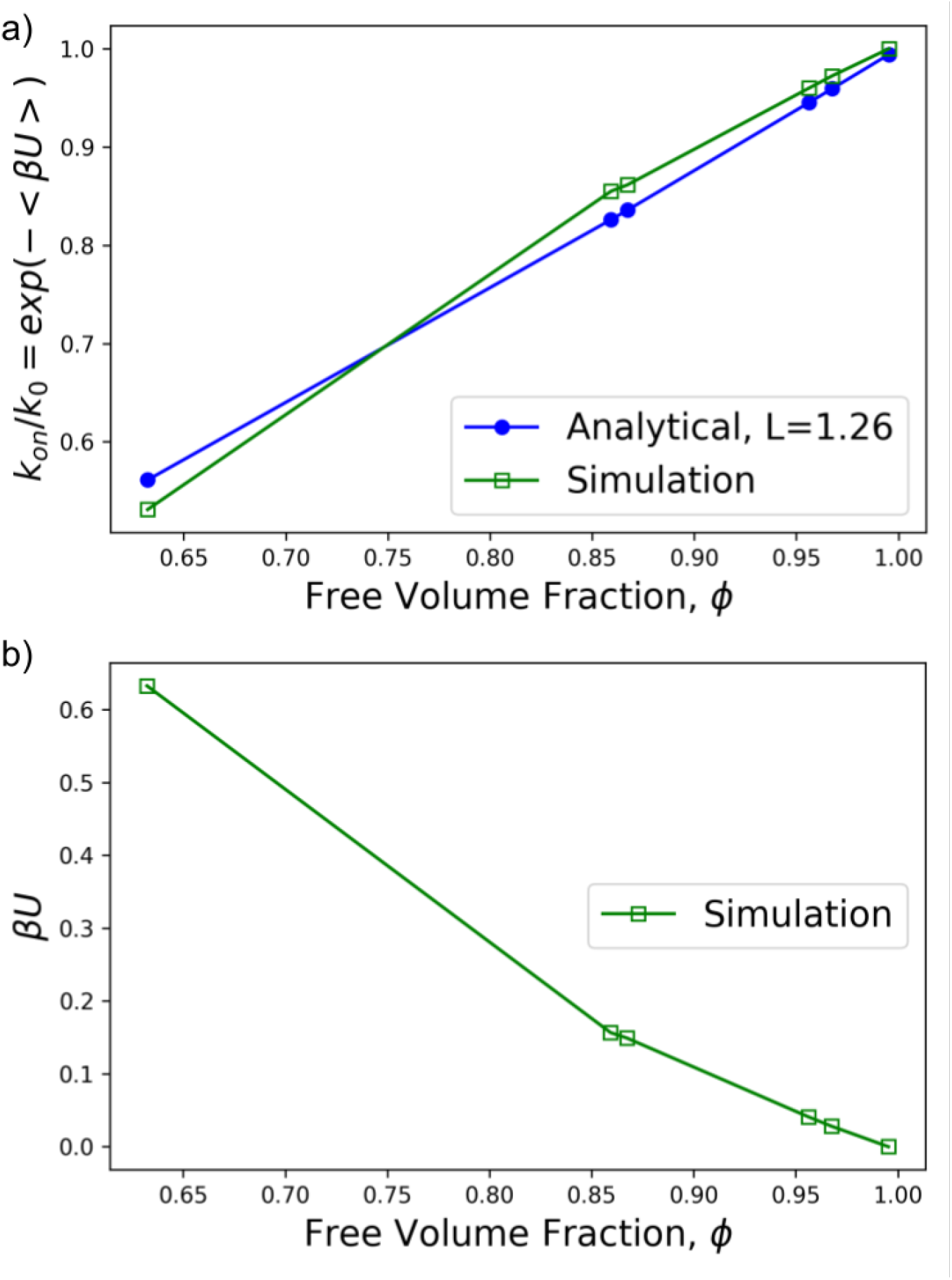
Effect of crowder density on normalized reaction rate coefficient. In the simulation data (green), the free volume fraction within the junction was varied by changing the number of non-reactive spherical inclusions (crowders). The analytical result (blue) is obtained from equation Eq. 6, with *L* obtained by fitting to the simulation data points. The agreement between the two curves demonstrates that the form of the dependence on *ϕ* is adequately described by Eq. 6.

Since inert crowders do not have direct enthalpic interactions with the substrate, we attributed the repulsive potential in Fig. 5 to increasingly unfavorable entropic interactions. To quantify this effect, we assumed that the diffusional volume comprises a large number of ‘sub-volumes’. In the absence of crowders, all sub-volumes are equally accessible and thus the configurational entropy of the substrate within the pore is maximized. When crowders were introduced, the number of accessible sub-volumes was reduced, which corresponded to a loss of entropy. We approximated these entropic contributions following arguments from Garcia *et al* [48], for which we define a configurational potential based on a substrate that can occupy N sub-volumes within the pore. This configurational potential is derived in Sect. S.2 and yields the following expression:

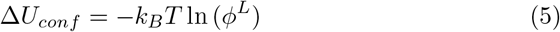

This expression relates the substrate’s accessible free volume to its configurational entropy, where L is a number related to the substrate concentration. Using this configurational energy in place of the electrostatic energy in Eq. 4 results in the following functional form for the dependence of the reaction rate coefficient on the free volume fraction, which validated against simulation data in Fig. 5.

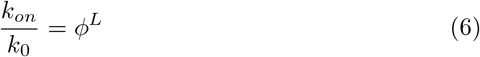

Because the reaction rate coefficients depend on the effective diffusion coefficient within the junction, Eq. 6 is similar to an empirical exponent sometimes used to express the dependence of the effective diffusion coefficient on the free volume fraction [49].

In Fig. 5 we compare the analytical result of Eq. 6 to the normalized reaction rate coefficients from finite element simulations, by selecting the value of *L* that best fits the simulation data. The agreement between the simulation data and the fitted analytical curve illustrates that the configurational potential energy of Eq. S11 is appropriate as a replacement for the electrostatic potential energy in Eq. 4 for the crowded condition. This suggests that the suppression of the ATP/CD39 association rate in the presence of crowders can be explained partly as a reduction in configurational entropy that the crowders impose.

With respect to diffusion rates in crowded media, attractive interactions with crowders can offset the configurational entropy of the substrate in the crowded domain, as suggested in Putzel *et al* [36]. To further investigate this effect, we present data in Fig. 6 examining the relationship between the Potential of Mean Force (PMF) and the electrostatic potential of the crowders, for different levels of crowding. The simulation results have an apparent linear relationship, with the slope depending on the free volume fraction within the junction. Thus, the PMF is generally proportional to the electrostatic potential of the crowders in the region investigated. Furthermore, the data illustrate a region of attractive electrostatic potentials where the resulting association rate is higher for more crowded systems than for less-crowded ones. This is evident, for example, at *V*_0_ =25 mV, where *U* = −0.23*k*_*b*_*T* for *ϕ* = 0.85 vs *U* = 0.0*k*_*b*_*T* for *ϕ* = 0.97. These data indicate the loss of configurational entropy due to crowders that k_*on,ATP*_ can be partially offset through introducing modest attraction between ATP and crowders. Crowders with an attractive potential will also compete with the enzyme for substrate. This effect would be expected to reduce the association rates for higher electrostatic potentials than are investigated in Fig. 6. Higher electrostatic potentials were investigated for a two-dimensional geometry in Fig. S1, where a minimum value of the PMF was indeed found. This minimum was generally in the range of *V*_0_ = 50 to 150 mV, depending on the free volume fraction within the junction. As the electric potential is increased beyond this range, the effective potential is increasingly thermodynamically unfavorable. As a result, ATP diffusion is slowed and manifests a reduced reaction rate. This mechanism of hindered diffusion is consistent with observations of diffusion over a ‘rough’ surface potential [50] that decreases the substrate diffusion coefficient. In this case, the roughness of the potential is created by repeated rising and falling of the potential that would be experienced on any path through the junction space between the crowders, where each crowder is a source for the potential.

**Figure 6:**
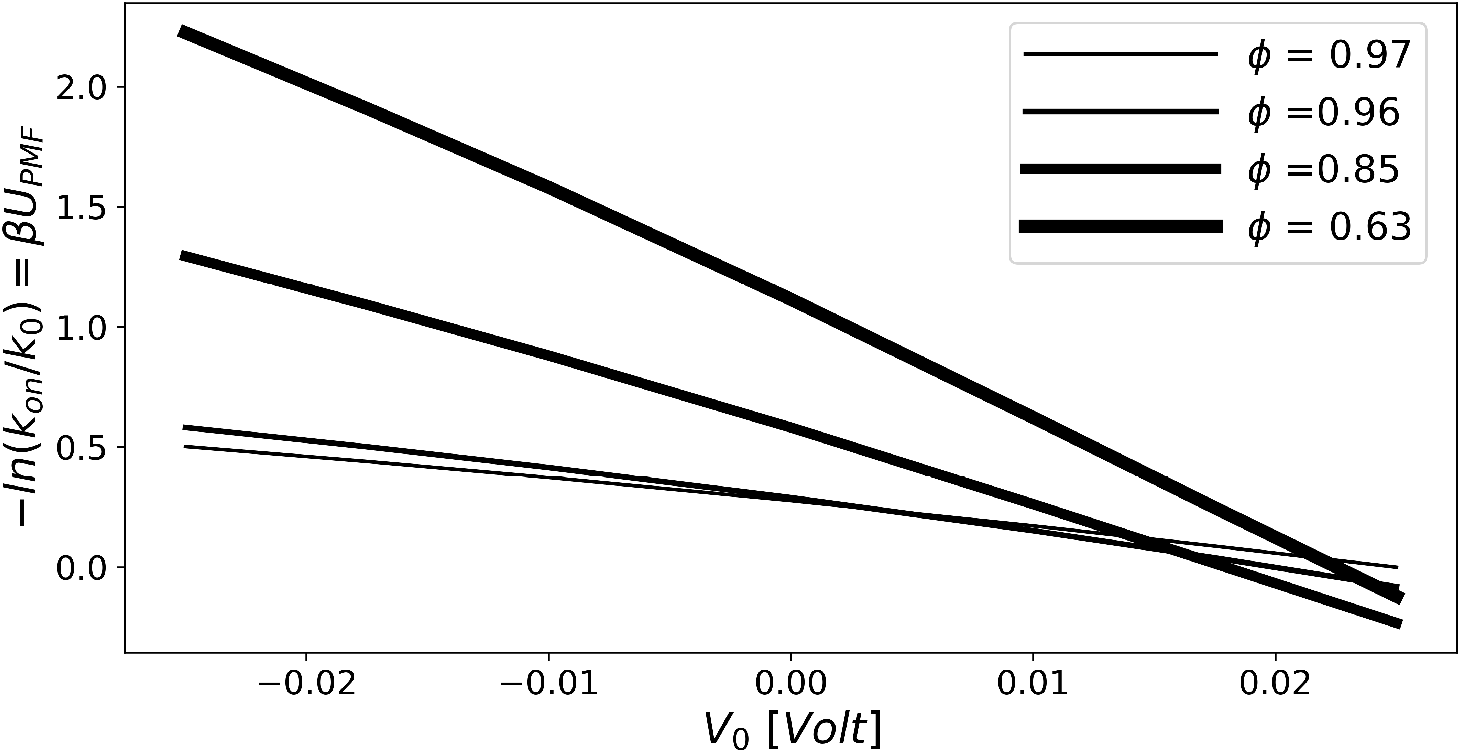
Effect of the electrical potential of nonreactive inclusions on the mean field potential. The vertical axis is the natural log of the normalized association rate coefficient from three-dimensional numerical simulations, which represents the mean force potential (see Eq. 4). The association rates are normalized to the result for *ϕ* = 0.97 with a crowder potential of 25 mV. A lower *U*_*PMF*_ value indicates higher rates of association. The horizontal axis is the electric potential boundary condition assigned to the crowders in the simulations, with positive values attractive to ATP and negative values repulsive. For some attractive potentials, the association rate can be higher for more crowded systems than for less crowded ones.

### 3.4 Crowders influence concerted nucleotidase activity in the synaptic junction

Our previous studies suggest that the kinetics of adenosine formation from concerted CD39 and CD73 activity are largely determined by the ATP association rate to CD39. However, the efficiency of adenosine formation, which we report as *κ*_*eff*_ =*k*_*ado*_/k_*on,ATP*_, is sensitive to factors including the CD39/CD73 separation distance. The rate of *k*_*ado*_, in particular, is influenced by the competition between the transport of AMP toward the CD73 enzyme versus its diffusion away from CD39. We therefore investigated how crowding modulates adenosine production efficiency, *κ*_*eff*_. First, we demonstrate for neutral crowders that *κ*_*eff*_ for the co-localized case was higher than when the enzymes were well-separated (15 nm)(See Fig. 7). In other words, *k*_*ado*_is higher when CD39 and CD73 are in close proximity. This was expected, as crowders distributed between the separated enzymes impeded AMP diffusion. As *ϕ* decreased, *κ*_*eff*_ decreased for the co-localized and separated enzymes. We subsequently evaluated *κ*_*eff*_ as a function of the crowders’ electric potential, which is summarized in Fig. 7. These data demonstrate that repulsive crowder/ATP interactions led to more efficient adenosine production, as those *κ*_*eff*_ values exceeded values from both attractive and neutral interactions. Attractive intereactions minimized *κ*_*eff*_, as the crowders competed with the enzymes for AMP by drawing the ATP away from CD73. Conversely, repulsive crowder interactions with AMP prevented substrate diffusion away from the reactive centers and thus promoted faster reaction rates at CD73. Hence, the crowders can tune the efficiency of concerted CD39/CD73 activity, depending on their density and long-range interactions. This behavior is analogous to bi-functional enzymes that utilize non-specific electrostatic interactions to channel intermediates toward reaction sites, as exemplified in dihydrofolate reductase-thymidylate synthase (DHFR-TS) [51–53]. [54]. Similar findings were reported for a crowded catalytic system by Roa *et al* [41].

**Figure 7:**
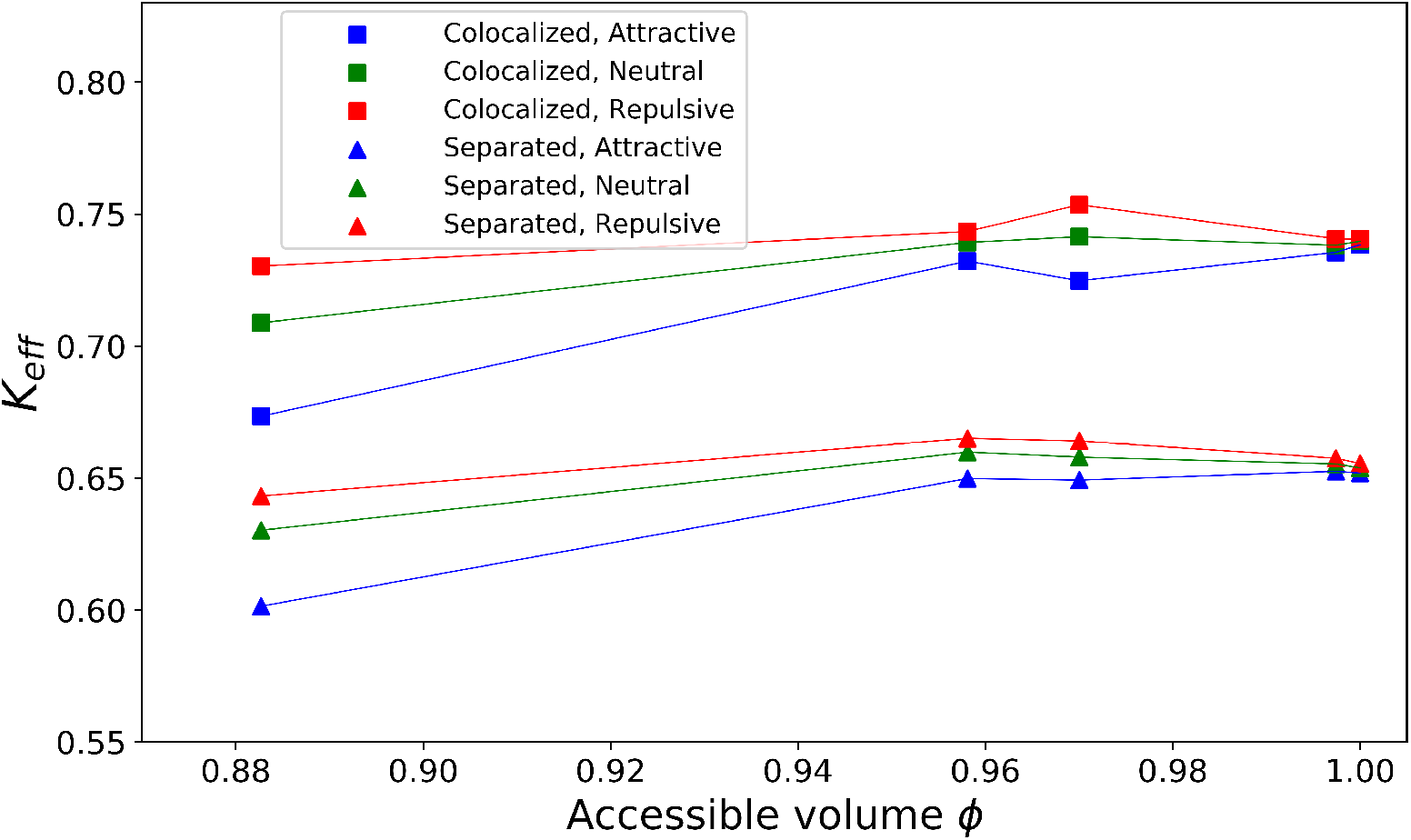
CD39/CD73 efficiency, K, with respect to the relative charged, non-reactive crowders. Results are shown for colocalized (square) and well-separated (triangle) CD39/CD73 enzymes. The magnitude of the electric potential for the attractive and repulsive crowders is 25mV.

Ultimately, the concerted activity of CD39/CD73 has the primary effect of controlling the composition of the nucleotide pool at the plasma membrane. The composition of this nucleotide pool determines a synapsed cell’s potential for agonizing receptors that sense extracellular ATP. We therefore report in Fig. 8 the steady state concentrations of ATP and AMP under crowded and crowder-free conditions. In agreement with our observations of *K*_*eff*_, crowded conditions generally impeded the entry of ATP and its metabolism in the junction, for all but attractive nucleotide/crowder interactions. The latter condition would prolong the stimulation of receptors that are sensitive to this nucleotide. We also include results for both colocalized enzymes (5 Å separation distance), as well as well-separated enzymes (150 Å separation distance). Under crowder free conditions, we recapitulated our earlier observations that ATP and AMP metabolism are higher for co-localized enzymes relative to well-separated reactive centers [1] as evidented by the comparatively lower ATP and AMP concentrations. Repulsive, and to a lesser extent, electrically neutral, crowders yield lower ATP and AMP concentrations, which is suggestive of enhanced nucleotide metabolism. Conversely, attractive nucleotide/crowder interactions yield the highest ATP and AMP concentrations. These data therefore suggest that while crowders enhance nucleotide pools at the plasma membrane owing to reducing CD39 activity and CD73 efficiency, the relative distribution of nucleotides is impacted by the density and charge of synaptic crowders.

**Figure 8:**
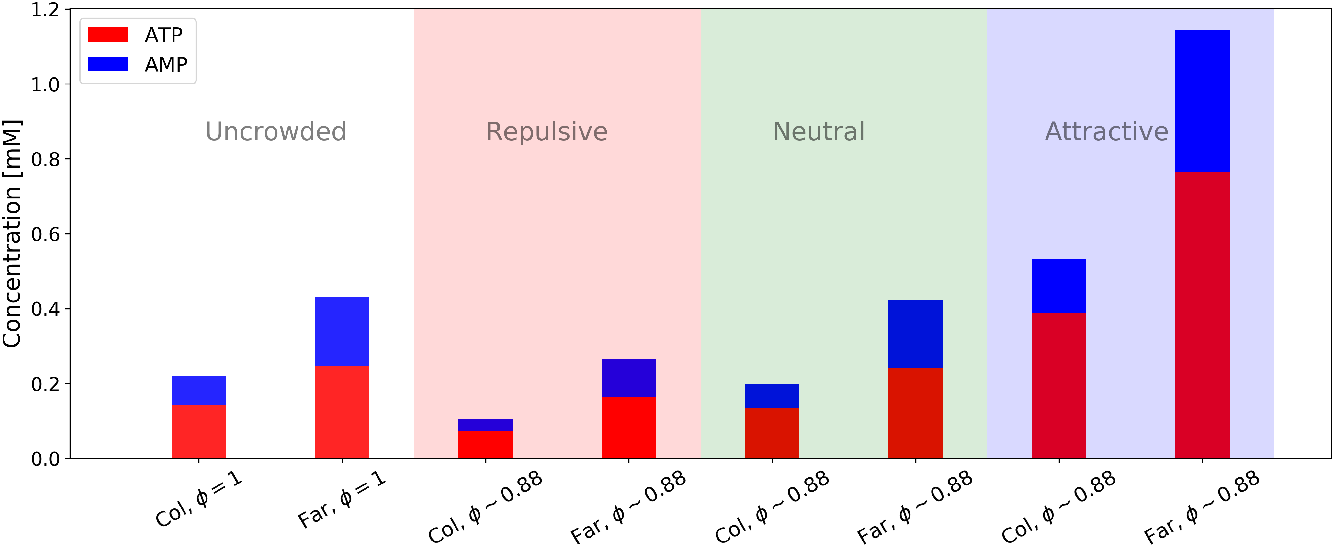
Concentration of ATP and AMP between the enzymes. “Col” refers to enzymes in close vicinity CD39/CD73; “Far” indicates enzymes separated by 15 nm.

### 3.5 Assumptions and Limitations

The assumptions related to the form of the configurational potential are described in the derivation within the Supplement. Many of the other assumptions involved in this work were also discussed previously [1]. Use of the linearized Poisson-Boltzmann equation for the electric potential requires relatively low ionic concentrations. We assumed that the reaction rates at the CD39 and CD73 enzymes were fast compared with the diffusion of nucleotides, such that the reactions were implemented as a reactive boundary condition. This assumption allows the solution of the steady-state diffusion equation rather than a time-domain equation. Enzymes and non-reactive inclusions are considered to be spherical instead of their actual surface geometry. However, a previous study demonstrated that protein structure has minor influence on the reaction rates [55]. Substrate molecules were modeled as point-particles and were assumed not to interact with one another. As a result, the reaction encounter distance between the substrate and an enzyme is the radius of the enzyme. Similarly, the geometry of the synaptic junction was simplified to a cylinder. The reservoirs at both ends of the junction had to be of finite size for numerical simulation, and a rectangular geometry was assumed. Simple concentration boundary conditions giving rise to a substrate concentration gradient along the junction were assumed. A single value was used for the externally imposed gradient, but other gradient values meeting the assumptions described above would not be expected to deliver qualitatively different results.

## 4 Conclusions

Our study provides quantitative insights into how crowders localized within synaptic junctions between closely-appositioned cells influence the kinetics of nucleotide metabolism by CD39 and CD73. In our prior work investigating coupled CD39/CD73 activity under confinement within synapses, we found that the overall production rate of adenosine from ATP and AMP was predominantly determined by the ATP/CD39 association rate. Therefore, we first demonstrate how crowders generally suppress k_*on,ATP*_ relative to crowder-free conditions, which could be rationalized in terms of a loss of configurational entropy to the ATP. Interestingly, attractive ATP/crowder interactions could counter this suppression to a small degree, in essence by providing a favorable thermodynamic offset to the loss of entropy. We additionally demonstrated that crowder/nucleotide interactions impact the efficiency of AMP conversion into adenosine. Specifically, repulsive AMP/crowder interactions restrict the diffusion of the nucleotide away from the CD39 and CD73 reactive sites, which improves AMP metabolism. Conversely, attractive AMP/crowder interactions tend to reduce reaction efficiency by drawing the substrate away from the ectonucleotidases. Lastly, we used this computational model to predict the relative distribution of the ATP, AMP and adenosine nucleotides next to the plasma membrane, where they could regulate membrane receptors. Our modeling results suggest that attractive interactions between the anionic nucleotides and crowders lead to higher steady state ATP and AMP concentrations at the membrane relative to crowder free conditions. These effects are mitigated when the CD39 and CD73 enzymes are co-localized and thereby reduce the impact of crowders on the diffusion rate of the AMP intermediate. Therefore, these computational results suggest that the interplay of CD39/CD73 activity, co-localization and the physiochemical properties of intrasynapse crowders help determine the pool of nucleotides available to stimulate synapse-bound membrane receptors.

## 5 Materials and methods

### 5.1 System

To analyze steady-state characteristics of nucleotide diffusion in a three-dimensional porous media, computer simulation similar to [1] is employed. The model, including junction, reservoirs, CD39 and CD73, is similar to previous work[1] and summarized in Fig. 9. We augmented our confined channel geometry with crowders of different densities and interaction strengths to determine their effects on CD39/CD73 reaction rates.

**Figure 9:**
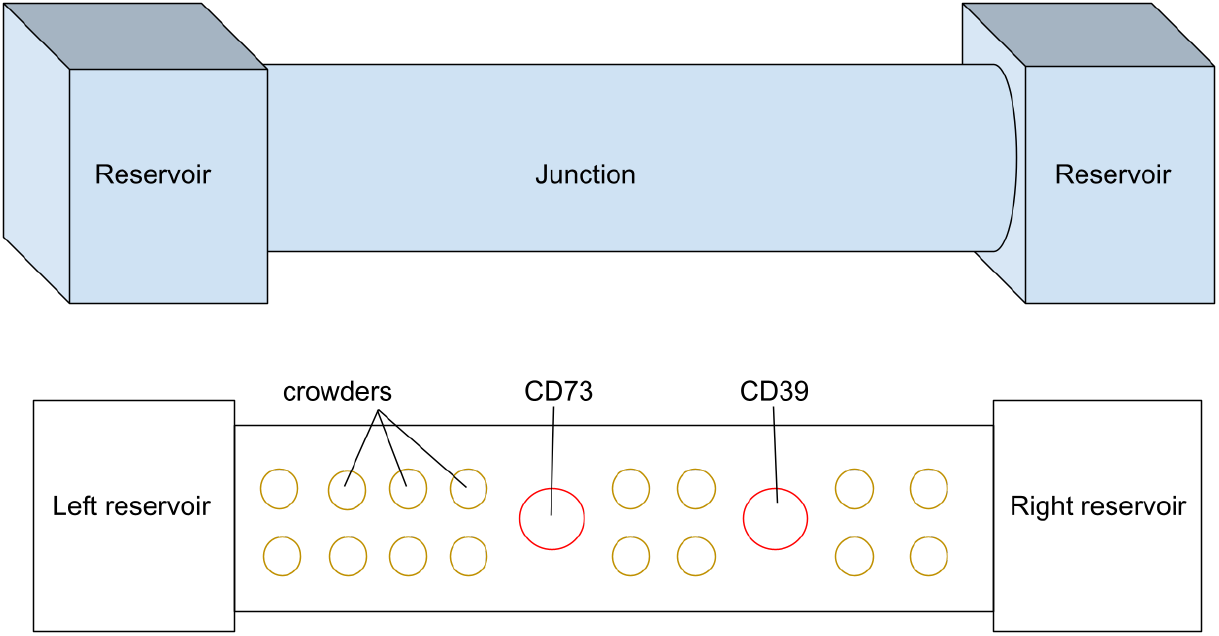
Sketch of model geometry. Top) Isometric view of the model including the reservoirs and the junction. Bottom) cross section of the system, showing the locations of coupled ectonucleotidases CD39/CD73 and non-reactive crowders around them.

Spherical inclusions on a regular lattice were incorporated into the model geometry by the following process. The channel was divided into two cylindrical sub-domains of equal size. This division was required to provide an internal surface for flux integral calculations. No spherical inclusion can cross this internal surface, as that would interfere with the flux calculation. Consequently, inclusions were placed separately within each sub-domain. The process of placing the inclusions added them sequentially. First, a set of points are located on a lattice with a specific distance from each other. The spherical inclusions are centered on these lattice points, and then checked for collision with each other as well as the sub-domain boundaries. A minimum separation distance was included in this check. If a collision was detected, the sphere was rejected, leaving an opening in the lattice. This process was repeated until the maximum number of inclusions was placed. The requirement that no sphere could overlap the sub-domain boundaries does lower the inclusion density near the boundaries. However, this effect should be limited to areas within one inclusion diameter of the boundaries.

### 5.2 Theory

The theoretical model of solving a steady-state partial differential equation is explained in [1]. In summary, the solution of the Fickian diffusion equation for a spherical surface in a bulk-like condition in the steady-state condition is

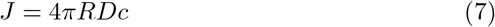

where *J* is the flux of species on the point of the surface, *R* is the radius of the spherical surface, D represents diffusion coefficient, and c is the nucleotide concentration.

In the presence of electrostatic potential,Φ, the Smoluchowski equation is required to be solved, which is given by:

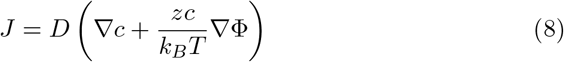

where *z* is the electric charge of the species, *k*_*B*_ is the Boltzmann constant, and *T* is the temperature. Under steady-state conditions and a continuity requirement, this equation can also be expressed as:

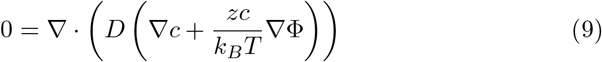

The electric potential in solution can be computed by solving the linearized Poisson-Boltzmann equation [56]:

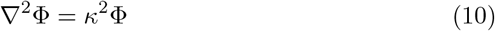

In which *κ* is the Debye-Hückel parameter, which is proportional to the square root of ionic strength *I*, and inversely proportional to the Debye-Hückel length *λ*.

In the study system, the electrostatic boundary conditions include setting the potential to zero at the reservoirs. The electrostatic potential on the surface of the crowders was variable.

We defined reaction rate *k*_*on*_ based on the total flux over the surface of the enzyme,

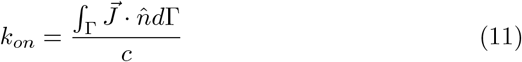

where *d*Γ is an element of surface area.

### 5.3 Numerical approach

Methodologies, the systems of partial differential equations, and boundary conditions are generally described in previous work [1]. We applied the Finite Element Method using the open-source finite element package FEniCS[57], version 2019.1.0, to conduct numerical solutions.

We executed Python-based analysis routines for setting up, solving, and post-processing of the finite element model. All of the processing code and data are publicly available at https://github.com/huskeypm/pkh-lab-analyses.

## 6 Supplement

### S.1 Figures

**Figure S1:**
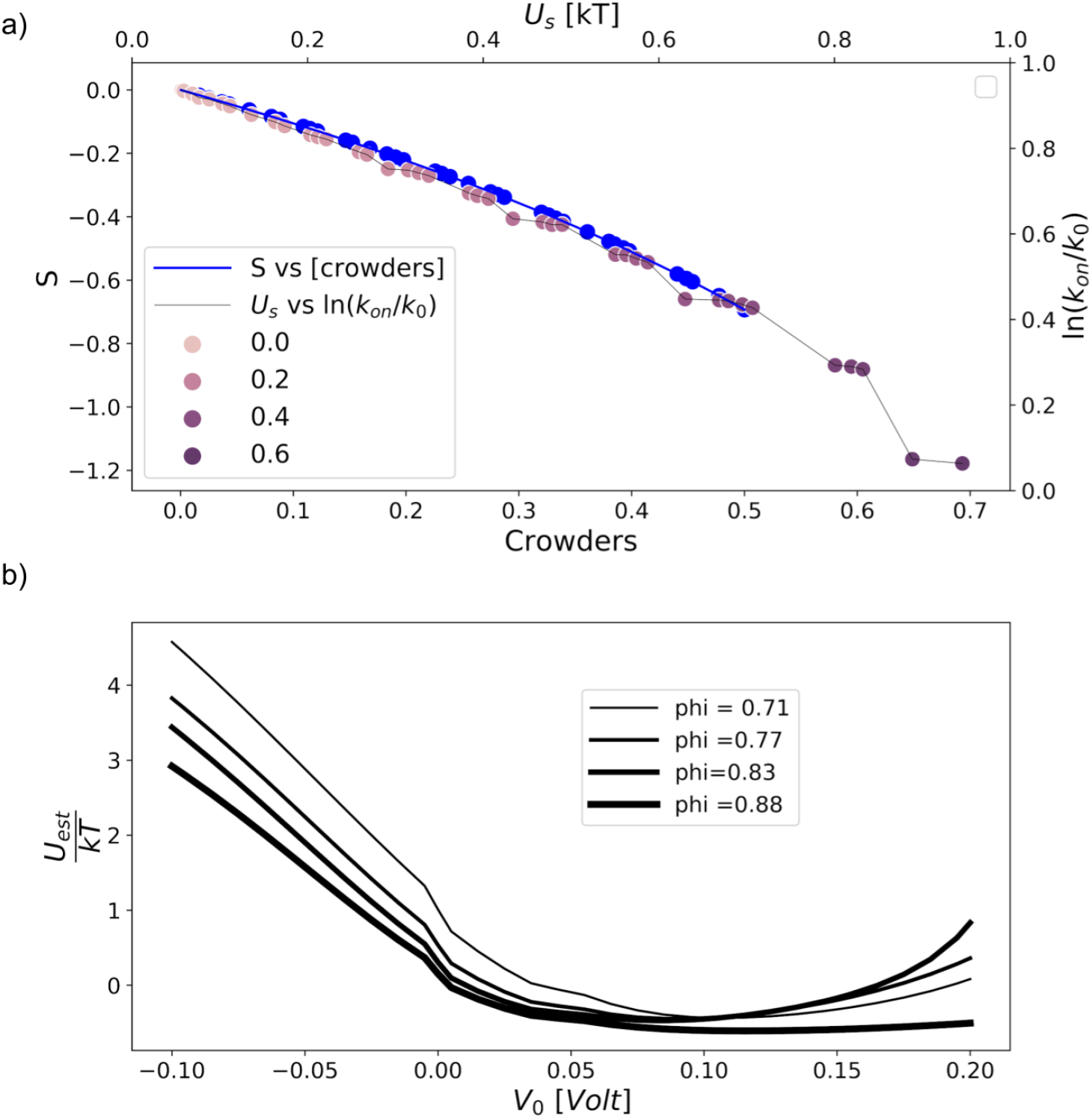
Configurational and electrostatic potential of the system as a function of crowders number in 2 dimensional. a) An analytical estimation of entropy and resulted in configurational potential based on that. b)Estimated interactional potential due to both configuration and electrostatic interactions. This is accomplished by representing the proteins as circles in a 2D plane. We have chosen the protein leucine-rich glioma inactivated 1 (LGI1) [42] to define the radius of these circles.

**Figure S2:**
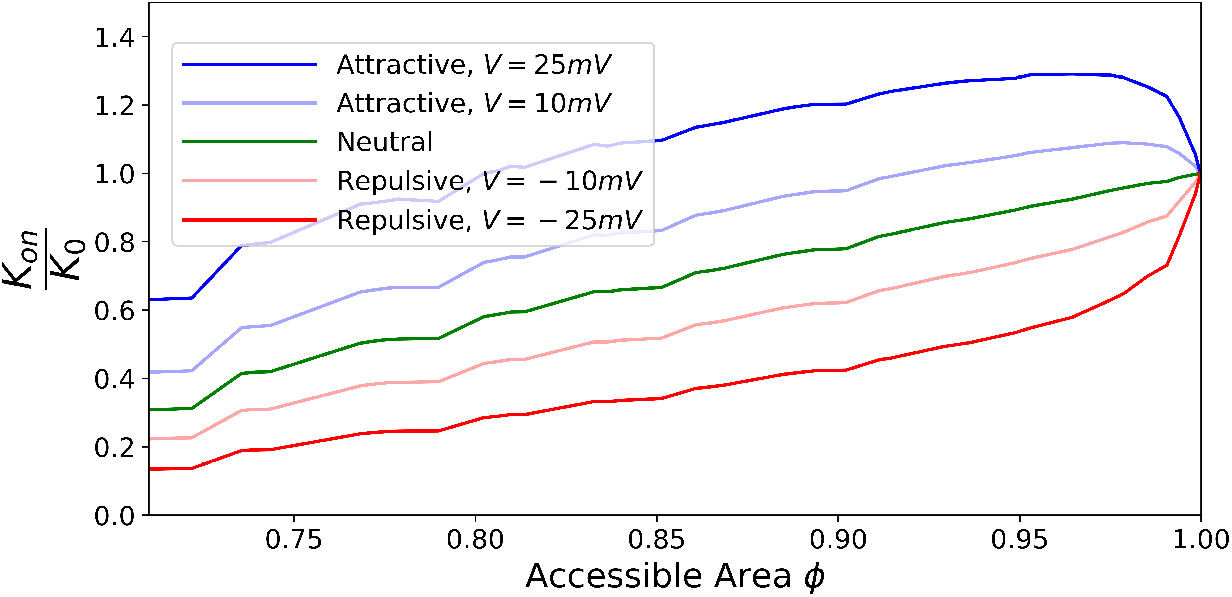
Effect of electrical potential of non-reactive inclusions on reaction rate, in two dimensions. See Fig. 3 for the corresponding results for three dimensions.

### S.2 Entropy in a crowded system

To analytically compute the configuration potential, we first obtain a relation for entropy as a function of accessible volume for the species. We then thermodynamically connect that to the potential, which leads to a relationship between the number of crowders and configurational potential. First, assume that the junction is divided into *N* subvolumes available to *L* ligands (substrate molecules), while *C* subvolumes are obstructed by crowders. This leaves *X* = *N* − *C* subvolumes available for ligands. From this point, the total number of microstates (*W*) of the configuration is equal to the number of possible ways to insert *L* indistinguishable ligands into *X* available subvolumes [58]:

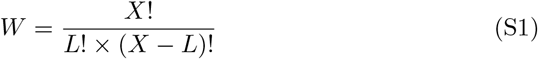

Since *L* ≪ *X*,

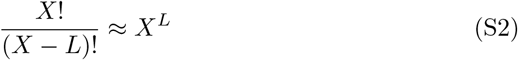

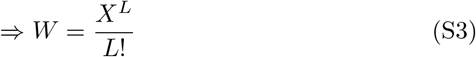

Applying the Stirling approximation to the logarithm of *W* leads to:

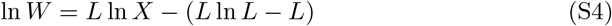

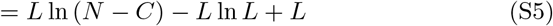

This leads to an explicit expression for the Boltzmann entropy:

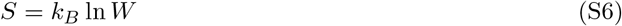

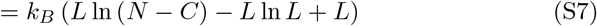

We define the configurational energy as the product of the configurational entropy and temperature.

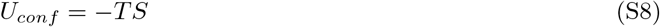

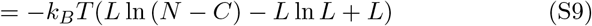

For the dilute condition in which *L* ≪ (*N* − *C*) this can be approximated as:

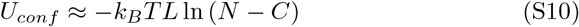

Which gives a relationship between configurational potential and the number of crowders. This can be represented based on 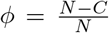, which defines the substrate-accessible volume:

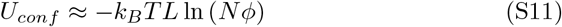

In the uncrowded case, *ϕ* = 1, and so the change in entropy from the uncrowded case to the crowded case is

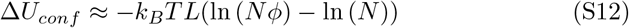

From the properties of logarithms, this simplifies to

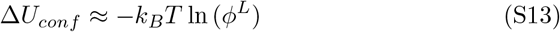

This approximate expression is used here as a general form, with the value of *L* as a fitting parameter. This fitting parameter would be expected to depend on factors not included in the foregoing analysis, such as the actual crowder shape and spatial distribution. The analysis suggests that this parameter may also depend on some measure of the substrate concentration.

## S Acknowledgements

Research reported in this publication, the release was supported by the Maximizing Investigators’ Research Award (MIRA) (R35) from the National Institute of General Medical Sciences (NIGMS) of the National Institutes of Health (NIH) under grant number R35GM124977. We thank Byeong Jae Chun for providing Fig. 1 based on renderings from Biorender.com.

